# Occurrence of Drosophilids in the Pernambuco’s coastal region (years 2000-2001) and the Frequency of chromosomal inversions found in *Drosophila malerkotliana*

**DOI:** 10.1101/2020.08.05.238634

**Authors:** Pierre Teodósio Félix, José Ferreira dos Santos

## Abstract

Drosophilids were collected in the *campus* of UFPE (Recife), Rio Doce (Olinda), Vila Velha (Itamaracá), Parque de dois Irmãos (Recife) and Charles Darwin Ecological Refuge (Igarassu), in the years 2000 and 2001, seeking to establish the frequency of occurrence of the various species of the genus Drosophila. In these collections, *D. malerkotliana* occurred in an average frequency of 64%, followed by *D. melanogaster*, with an average frequency of 23%. The frequency of *D. malerkotliana* was higher than that of *D. melanogaster* in localities with high and medium degree of urbanization. Despite the great distance from the distribution center of the species (Africa), *D. malerkotliana* presented high polymorphism of chromosomal inversions. In the locality of Rio Doce, inversions In(IIL)24B-39A and In(IIL)21-25, occurred with frequencies of 18.5% and 100%, respectively, while inversion In(IIIR)84-88B occurred with a frequency of 100%. In the population of *campus* UFPE (Recife), also with high urbanization, two inversions were found, In(IIIR)93C-94A and In(IIIL)64A-72A with frequencies of 37.1 and 60%, respectively. In the population of The Parque de Dois Irmãos (Recife), with medium urbanization, four inversions were visualized: In(IIIL)70C-73B (66%); In(IIIR)93C-94A (52%); In(IIL)24B-39A (33%); and In(IIR)50B-51A (33%). In the population of the Charles Darwin Ecological Refuge (Igarassu), of low urbanization, only the inversion In(IIIR)100A-98B(?) was found. −88(?), with 100% frequency. These data suggest that the polymorphism of inversions is higher in localities with greater urbanization, possibly due to the longer colonization time, which allowed the accumulation of genetic variations.

## Introduction

*Drosophila malerkotliana* is a species of sub cosmopolitan distribution that has been highlighted by its easy adaptation in the colonization of new environments, a high rate of occurrence and recent dispersion, as a consequence of human activity (LEMEUNIER *et al*., 1986). Originally from Africa, *D. malerkotliana* is currently colonizing Brazil in the North-South direction, having been registered in the country for the first time in the 1970s in the Northeast (VALE and SENE, 1980) and in the Amazon in the 1980s (MORAIS et al., 1995). The Southern limit of your most recently recorded distribution is Santa Catarina (from TONI, 1998). The high invasive power of this species is probably determining major ecological changes in the Brazilian fauna of Drosophila, by the displacement or substitution of native or previously established species (NASCIMENTO, 2000).

In regions of Latin America, such as the Caribbean, this species is already among the dominant ones in the *drosophilid* guild (MORAIS et al., 1995). In the State of Pernambuco, *D. malerkotliana* occurs at high relative frequencies until 1999, associated with a large number of paracentric inversions and at least one pericentric inversion complex (NASCIMENTO, 2000). In this work, the frequencies of drosophilid species were recorded, as well as the inversions that occur in the polythenic arms of *D. malerkotliana* of four localities of the coastal region of the State of Pernambuco were identified and quantified. In addition, a link was established between the chromosomal inversion polymorphism and the invasive potential of the species, based on the frequency of occurrence of the species in the Pernambuco coast.

## Material and Methods

### Species collection and identification

Drosophilids were collected in different locations in the coastal region of the State of Pernambuco, with different degrees of urbanization. The collection sites with a marked level of urbanization were two localities within the Greater Recife region, the UFPE *Campus*, within the City of Recife and Rio Doce, a neighborhood in the municipality of Olinda. In these localities there is a large number of buildings, without preserved native vegetation. From the localities with medium level of urbanization were chosen the Parque de dois irmãos, which even being a forest reserve with good preservation of native vegetation (secondary forest spot), is located within the city of Recife and surrounded by environments with high levels of urbanization. The locations chosen with a low level of urbanization were the Charles Darwin Ecological Refuge, a reserve of secondary Atlantic forest, well preserved in recent decades, located in the municipality of Igarassu, and Vila Velha, in Itamaracá (Figure 1). The samples were collected during the years 2000 and 2001, following the availability of adult individuals, which varies according to climatic conditions.

The collections were made using banana baits, taken to the field and distributed in cylindrical glass tubes with a capacity of 150ml, or by collecting eggs and larvae in fruits in fermentation. A stereoscopic microscope was used for taxonomic observation and identification of adult individuals, according to identification key (FREIRE-MAIA and PAVAN, 1949). For differentiation between *D. melanogaster* and *D. malerkotliana*, the absence of tarsal comb was considered in males of the latter species (VAL and SENE, 1980).

**Figure 1.**
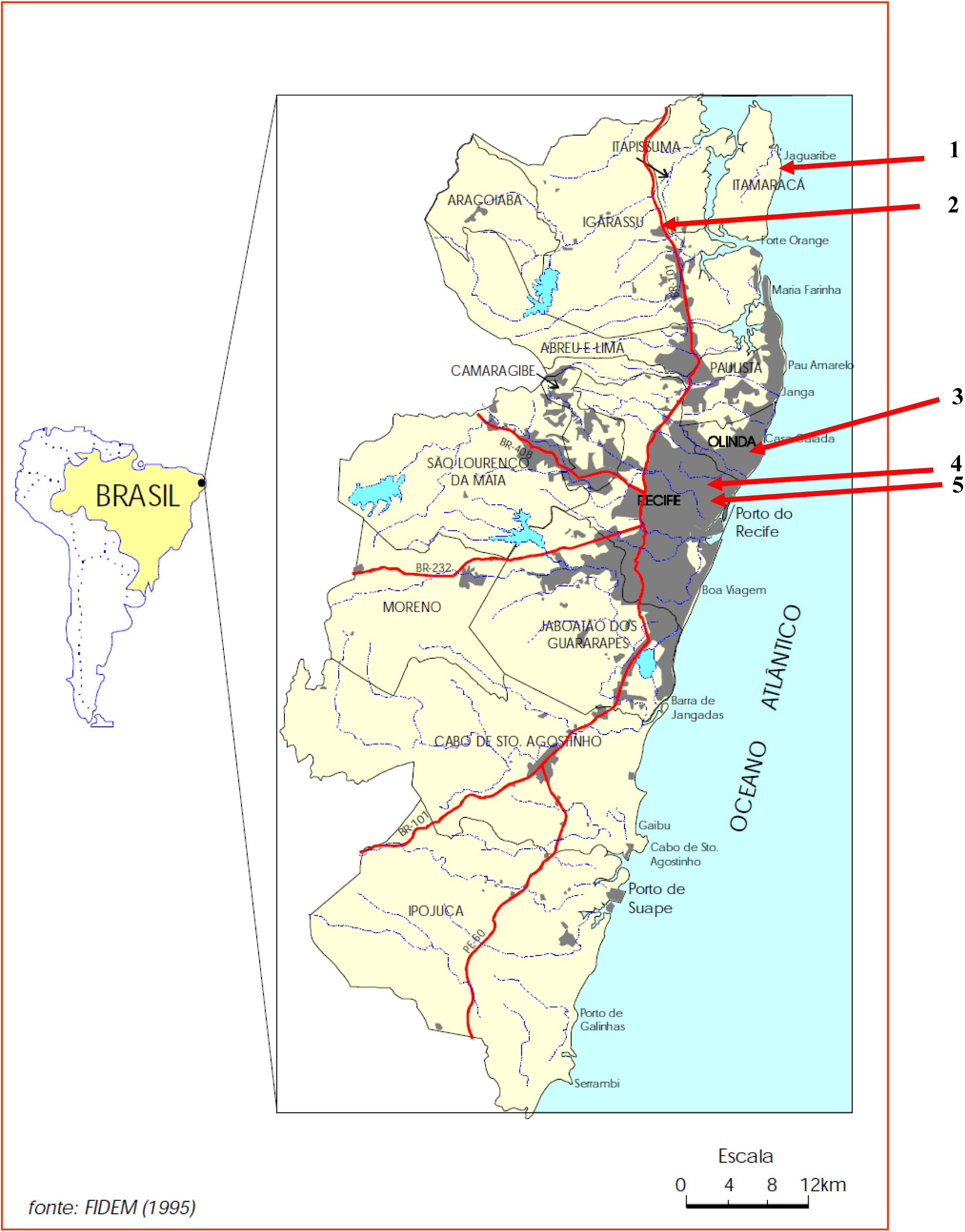
Map of the Coastal Region of Pernambuco, indicating the collection sites: Vila Velha in Itamaracá (1); Charles Darwin Ecological Refuge in Igarassu (2); Rio Doce in Olinda (3); UFPE *Campus* (4) and Parque de dois irmãos in Recife (5). Map adapted by FIDEM (1995).

### Cytological preparations

The females of *D*.*malerkotliana* born from each population were individually peaked to establish iso-females in glass containing culture medium prepared with corn flour, fresh organic yeast and banana, being daily peaked and fed with yeast. Sixty-five iso-females strains from campus UFPE, 23 from Rio Doce, 12 from parque de dois irmãos and 19 from Charles Darwin Ecological Refuge were analyzed.

The slides of *D. malerkotliana* polythenic chromosomes were prepared by crushing the dissected salivary glands of third stage larvae, fixed with a solution of acetic acid, water and lactic acid, in a 3:2:1 ratio. The material was stained with 1% aceto lactic orcein (1 g of orcein, 45 ml of acetic acid, 25 ml of lactic acid 85% and 30 ml of distilled water), according to ASHBURNER (1967). After the best points were scored, the chromosomes were photographed under Leica Microscope in increase of 100 × 1.25 with Kodak’s TMAX Iso 100 film. The breaking points of the inversions were established by comparison with the photomap made by FÉLIX *et al*. (2002).

## Results

In the period studied *D. malerkotliana* was the most represented species, reaching 66% in 2000 and 61% in 2001, followed by *D. melanogaster*, with frequencies of 25 and 21% in those same years. The frequency, around 10%, also draws attention to the frequency of another invasive drosophilid, *Zaprionus indianus* (Figure 2).

**Figure 2.**
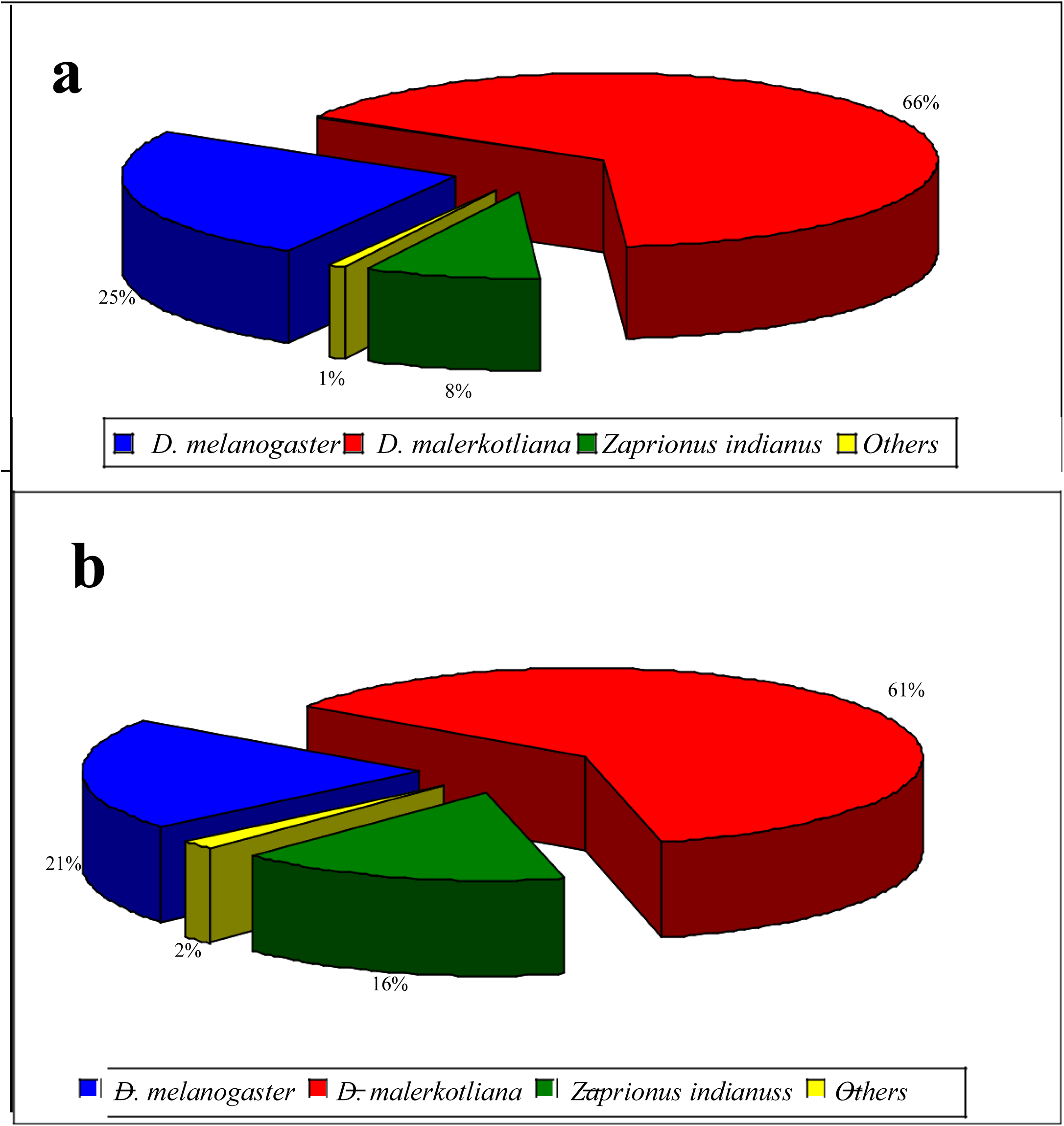
Frequency of drosophilids collected in the years 2000 (a) and 2001 (b) in the coastal region of Pernambuco, as shown in the map in Figure 1. Note the high frequencies of the invasive species *D. malerkotliana* and *Zaprionus indianus*.

Both in 2000 and 2001, the frequency of occurrence of *D. malerkotliana* was higher than that of D. melanogaster in the locations with high urbanization (UFPE *Campus*, Rio Doce) and medium (Parque Dois Irmãos), and the opposite is observed in the low urbanization locations (Vila Velha, Charles Darwin Ecological Refuge), as can be seen in Table 1.

**Table 1.**
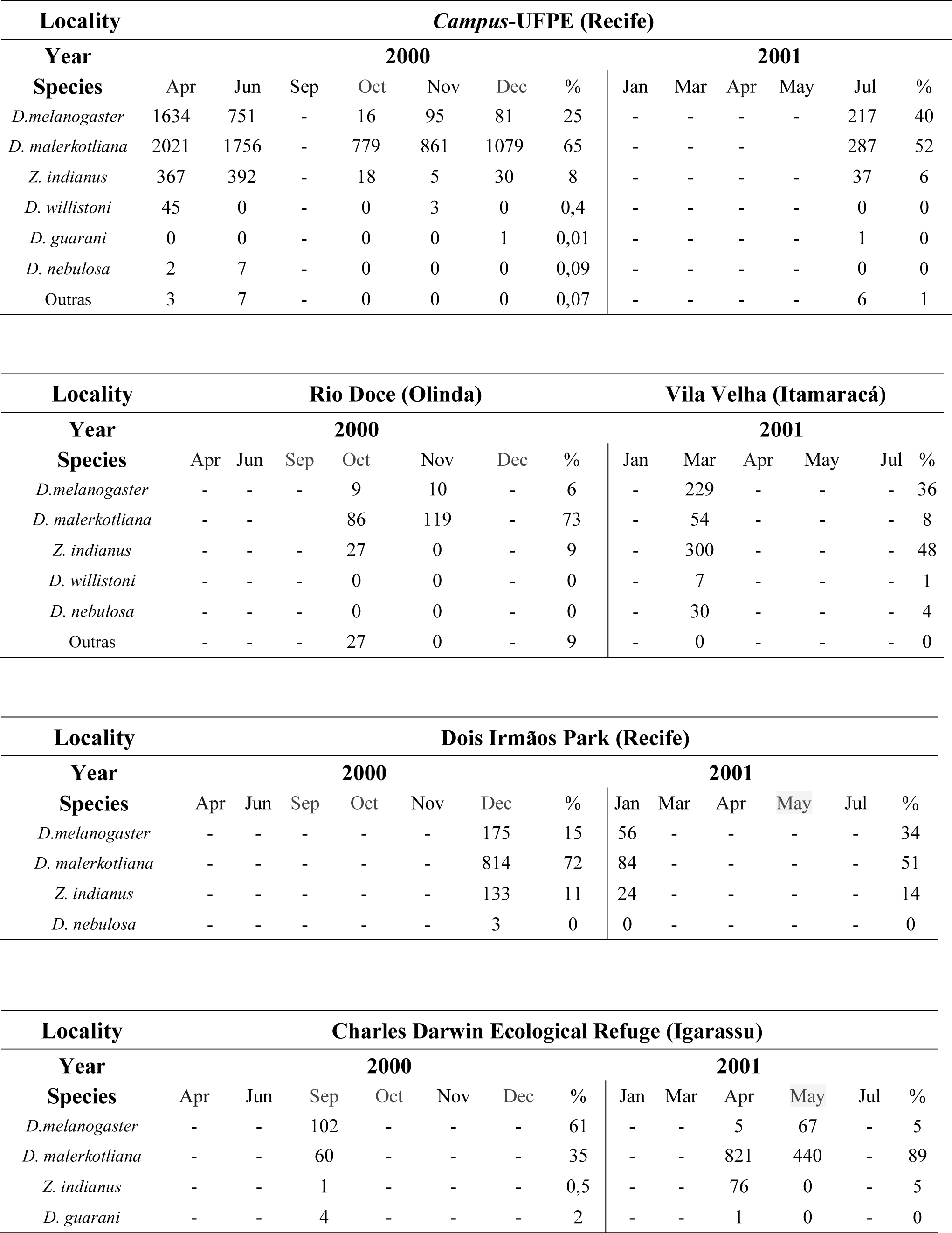
Frequency of Drosophilids occurrence in the in the Pernambuco’s coastal region (years 2000-2001).

| Locality |  | Campus-UFPE (Recife) |  |  |  |  |  |  |  |  |  |  |  |
| --- | --- | --- | --- | --- | --- | --- | --- | --- | --- | --- | --- | --- | --- |
| Year |  | 2000 |  |  |  |  |  | 2001 |  |  |  |  |  |
| Species | Apr | Jun | Sep | Oct | Nov | Dec | % | Jan | Mar | Apr | May | Jul | % |
| *D.melanogaster* | 1634 | 751 | - | 16 | 95 | 81 | 25 | - | - | - | - | 217 | 40 |
| *D. malerkotliana* | 2021 | 1756 | - | 779 | 861 | 1079 | 65 | - | - | - |  | 287 | 52 |
| *Z. indianus* | 367 | 392 | - | 18 | 5 | 30 | 8 | - | - | - | - | 37 | 6 |
| *D. willistoni* | 45 | 0 | - | 0 | 3 | 0 | 0,4 | - | - | - | - | 0 | 0 |
| *D. guarani* | 0 | 0 | - | 0 | 0 | 1 | 0,01 | - | - | - | - | 1 | 0 |
| *D. nebulosa* | 2 | 7 | - | 0 | 0 | 0 | 0,09 | - | - | - | - | 0 | 0 |
| Outras | 3 | 7 | - | 0 | 0 | 0 | 0,07 | - | - | - | - | 6 | 1 |

| Locality |  | Rio Doce (Olinda) |  |  |  |  |  | Vila Velha (Itamaracá) |  |  |  |  |  |
| --- | --- | --- | --- | --- | --- | --- | --- | --- | --- | --- | --- | --- | --- |
| Year |  | 2000 |  |  |  |  |  | 2001 |  |  |  |  |  |
| Species | Apr | Jun | Sep | Oct | Nov | Dec | % | Jan | Mar | Apr | May | Jul | % |
| *D.melanogaster* | - | - | - | 9 | 10 | - | 6 | - | 229 | - | - | - | 36 |
| *D. malerkotliana* | - | - |  | 86 | 119 | - | 73 | - | 54 | - | - | - | 8 |
| *Z. indianus* | - | - | - | 27 | 0 | - | 9 | - | 300 | - | - | - | 48 |
| *D. willistoni* | - | - | - | 0 | 0 | - | 0 | - | 7 | - | - | - | 1 |
| *D. nebulosa* | - | - | - | 0 | 0 | - | 0 | - | 30 | - | - | - | 4 |
| Outras | - | - | - | 27 | 0 | - | 9 | - | 0 | - | - | - | 0 |

| Locality |  | Dois Irmãos Park (Recife) |  |  |  |  |  |  |  |  |  |  |  |
| --- | --- | --- | --- | --- | --- | --- | --- | --- | --- | --- | --- | --- | --- |
| Year |  | 2000 |  |  |  |  |  | 2001 |  |  |  |  |  |
| Species | Apr | Jun | Sep | Oct | Nov | Dec | % | Jan | Mar | Apr | May | Jul | % |
| *D.melanogaster* | - | - | - | - | - | 175 | 15 | 56 | - | - | - | - | 34 |
| *D. malerkotliana* | - | - | - | - | - | 814 | 72 | 84 | - | - | - | - | 51 |
| *Z. indianus* | - | - | - | - | - | 133 | 11 | 24 | - | - | - | - | 14 |
| *D. nebulosa* | - | - | - | - | - | 3 | 0 | 0 | - | - | - | - | 0 |

| Locality |  | Charles Darwin Ecological Refuge (Igarassu) |  |  |  |  |  |  |  |  |  |  |  |
| --- | --- | --- | --- | --- | --- | --- | --- | --- | --- | --- | --- | --- | --- |
| Year |  | 2000 |  |  |  |  |  | 2001 |  |  |  |  |  |
| Species | Apr | Jun | Sep | Oct | Nov | Dec | % | Jan | Mar | Apr | May | Jul | % |
| *D.melanogaster* | - | - | 102 | - | - | - | 61 | - | - | 5 | 67 | - | 5 |
| *D. malerkotliana* | - | - | 60 | - | - | - | 35 | - | - | 821 | 440 | - | 89 |
| *Z. indianus* | - | - | 1 | - | - | - | 0,5 | - | - | 76 | 0 | - | 5 |
| *D. guarani* | - | - | 4 | - | - | - | 2 | - | - | 1 | 0 | - | 0 |

The frequencies of chromosomal inversions of the populations of UFPE *campus*, Rio Doce, Parque de dois Irmãos and Charles Darwin Ecological Refuge were determined (Table 2). In the population of the UFPE *campus*, two inversions were observed, one in the IIIR arm, In(IIIR)93C-94A with a frequency of 37.1%, and one in the IIIL arm, In(IIIL)64A-72A with a frequency of 60%. In the population of Rio Doce, three inversions were identified, two of them in the IIL arm, In(IIL)24B-39A with a frequency of 18.5% and In(IIL)21-25 with 100%, and one in the IIIR arm, In(IIIR)84-88B, with a frequency of 100%. In the population of The Parque de dois Irmãos, four inversions were found, one of them in the IIIL arm, In(3L)70C-73B with frequency of 66%, another in the IIIR arm, In(3R)93C-94A with frequency of 52%, another in the IIL arm, In(2L)24B-39A with frequency of 33% and one in the IIR arm, In(2R)50B-51A with frequency of 33%. In the population of the Charles Darwin Ecological Refuge, a single inversion was found in the IIIR arm, In(3R)100A-98B? −88? 100% frequency.

**Table 2.**
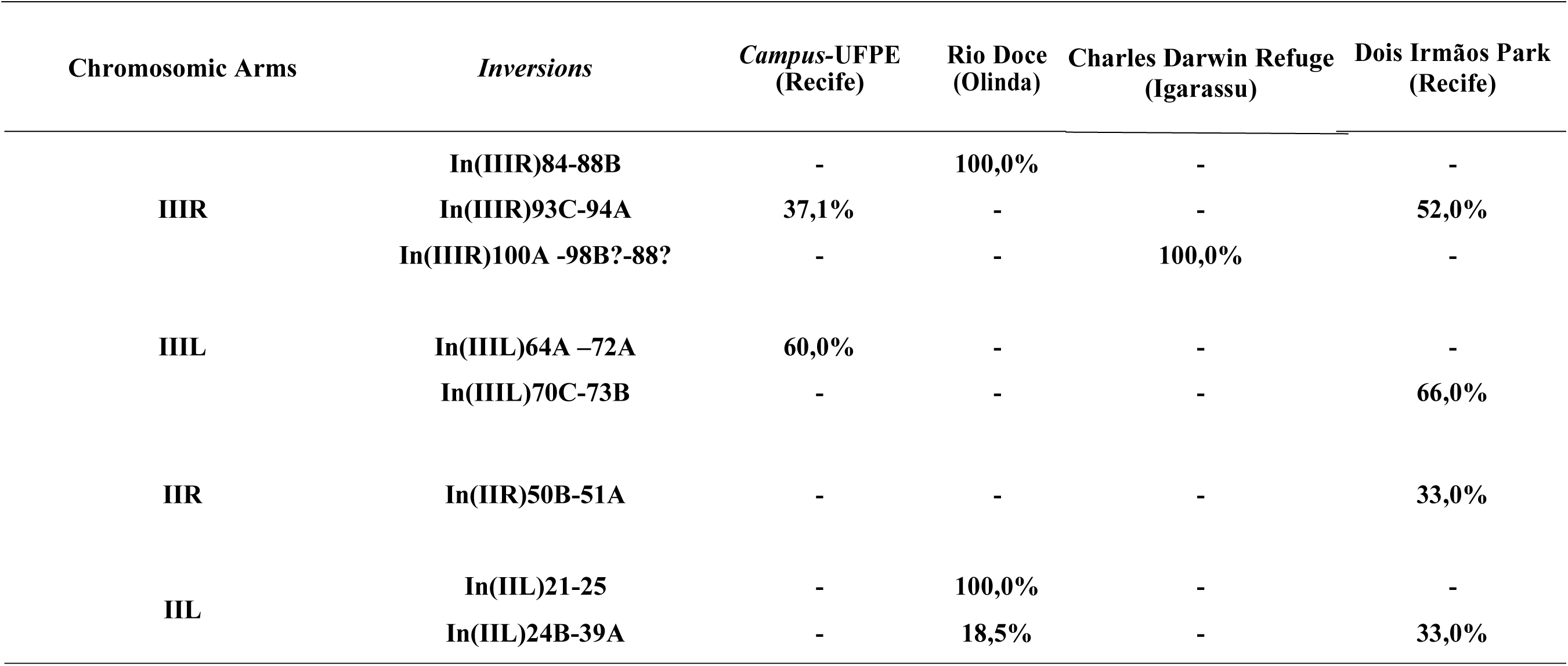
Frequencies of paracentric inversions in heterozygosis in the polythenic chromosomes of populations of *D. malerkotliana* from the Pernambuco’s coastal region. The breakpoints of the inversions were determined based on the photomap made by FÉLIX *et al* (2002).

## Discussion

The species of Drosophilids from northeastern Brazil are scarcely studied, and only some species of the **melanogaster** and **willistoni** group are recorded (VAL and SENE, 1980; SENE *et al*., 1980; EHRMAN and POWELL, 1982). In this work, populations from five localities in the coastal region of Pernambuco were studied, seeking to relate the drosophilid fauna to the different degrees of environmental urbanization, as verified between the urban area of Porto Alegre and the area without urbanization in Rio Grande do Sul (SANTOS and VALENTE, 1990; BONORINO *et al*., 1993; ROHDE and VALENTE, 1996; VALIATE and VALENTE, 1996, 1997).

The largest variety of species was found in the Charles Darwin Refuge, probably due to the better ecological conditions and their relative distance (approximately 7 km) from large urban centers. However, the species *D. malerkotliana* and *D. melanogaster* were more frequent in environments with medium (Parque de Dois Irmãos) and high urbanization levels (UFPE Campus and Rio Doce). Possibly, like *D. melanogaster, D. malerkotliana* is also preferentially distributed in a way associated with human activities (LEMEUNIER *et al*. 1986).

Due to its potential invader *D. malerkotliana* now stands out as the predominant species in the coastal region of the State of Pernambuco. Confirming the extensive alteration of the current fauna in relation to that described in the literature (VAL and SENE, 1980; SENE *et al*., 1980; VAL *et al*., 1981; EHRMAN and POWELL, 1982), found at relatively high frequency (10%) the drosophilid *Zaprionus indianus*, whose presence in the State was initially recorded in 2000 (FÉLIX et al., 2001). Other species such as *Drosophila kikawai, D. nebula* and *D. guarani* had frequencies lower than 1% in the years 2000 and 2001. Nevertheless, it was not expected to find the latter in the Northeast, because it is a species considered cold climate (VAL *et al*., 1981).

Numerous factors such as limitation of feeding resources and low rainfall rates may have caused changes in the drosophilid fauna in northeastern Brazil. In addition to these factors, competition between species contributed to the replacement of *D. melanogaster* by *D. malerkotliana* in the Northern Region (MARTINS, 1995; SANJAD, 1997) and Nordeste (NASCIMENTO, 2000) of Brazil, as also observed by VALENTE *et al*. (1993) and VALIATE and VALENTE (1996) for *D. willistoni* in Rio Grande do Sul. These changes in the Drosophilids fauna may have also led to important qualitative changes in the composition of the communities of other associated organisms, such as yeasts, which have a direct association with drosophilids for their dispersion (BEGON, 1982). Due to the lack of previous studies, the existing relationships derived from these changes cannot be fully evaluated.

The colonization of new environments by Drosophilids is a common phenomenon, and species substitution has been observed with a certain frequency. The invasive potential of the species and the ease of adaptation to austere environments has been attributed, by several authors, to the loss or gain of variability by selection forces. VALIATE and VALENTE (1997) observed a substitution of *D. willistoni* by *D. paulistorum* in the city of Porto Alegre, demonstrating that *D. paulistorum* has a large number of chromosomal inversions, thus increasing its variability and adaptive power. The gene arrangements contained in the inversion loops are inherited in a simple Mendelian way, passing to the descendants as a single gene (TAYLOR and POWELL, 1983). The variation in polymorphism results from the ability of species to explore different environments and plays an adaptive role for various aspects related to the survival of individuals and reproduction (SPERLICH and PFRIEM, 1986).

From the genetic point of view, inversion has as a consequence the restriction, or even total blockage of gene recombination in the inverted region. When pairing is inverted or a-synaptic, chiasm usually does not occur in this region and, when it occurs, forms chromatids with deletions and duplications, due to incorrect pairing. Evolutionarily this blockage causes some heterozygotes for inversions to be more adapt than their homozygous forms, because only the genes contained in the inversion are maintained in heterozygosis (TAYLOR and POWELL, 1983). This hypothesis is corroborated by the fact that in some populations there is a certain inversion in 100% of the individuals analyzed, such as In(3R)84-88B in the population of Rio Doce (Olinda) and In(3R)100A-98B? −88? population of the Charles Darwin Ecological Refuge (Table 2).

The populations of *D. malerkoltiana* present a high degree of polymorphism of chromosomal inversions, even though it is a species that recently invaded Brazilian territory (VAL and SENE, 1980). Associated with the polymorphism of male tergite coloring (NASCIMENTO, 2000), inversion polymorphism characterizes a genetic variability that can enable the species to colonize this type of colonization, since the most polymorphic populations tend to occupy large areas because they have a variety of gene arrangements that allow the occupation of a wide variety of habitats (TAYLOR and POWELL, 1983). It is worth remembering that due to the similarity of the banding pattern between the sections of the polythenic chromosomes of *D. malerkotliana*, it was not possible to quantify the frequencies of homozygotes for the “standard” or “inverted” arrangements.

It was found that in the low-level urbanization environment, the Charles Darwin Ecological Refuge in Igarassu, the population did not present high polymorphism of chromosomal inversions. However, the inversion In(3R)100A-98B? −88? it was found in all the polythenic nuclei observed, showing conservation in the karyotypic constitution, possibly due to the recent establishment of the population. The populations of localities with a high level of urbanization, such as those of the UFPE *Campus* and Rio Doce, presented together five chromosomal inversions, present in all 4 autosomal arms. However, one of the inversions, In(2L)24B-39A, is common to the population of Parque Dois Irmãos, which is an environment with an average level of urbanization. Based on these observations, it is suggested that there was a decrease in the polymorphism of the chromosomal inversions of the populations, in relation to the increase in the level of urbanization of the localities.

## Attachments

**Figure 2.**
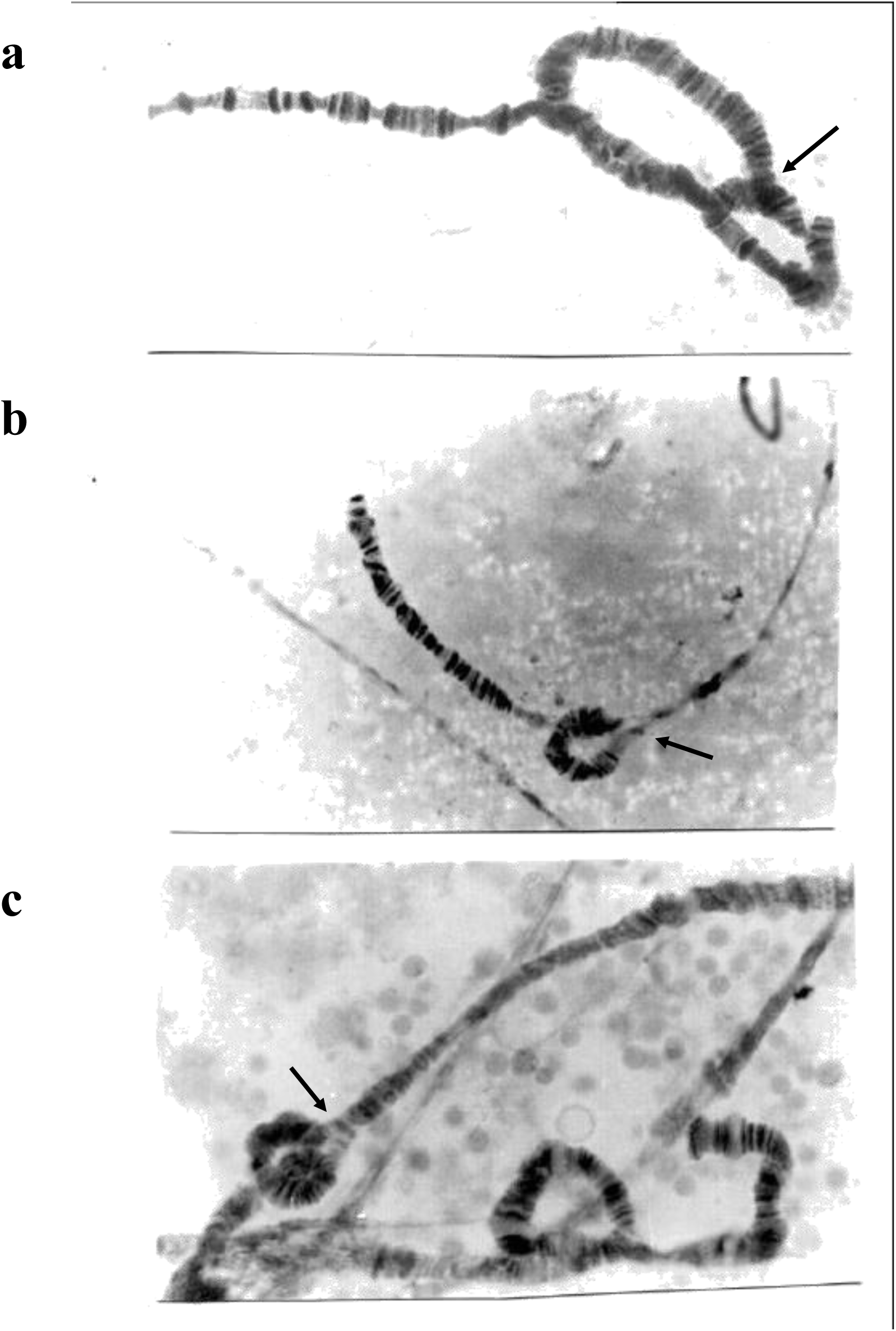
Chromosomal inversions in the IIIR arm of *D. malerkotliana*. In(IIIR)100A-98B?-88?, with a frequency of 100% of the population of the Charles Darwin Ecological Refuge (a). In(IIIR)93C-94A, with frequency of 37.1% of the population of Campus-UFPE (b). In(IIIR)84-88B, with frequency of 100% of the population of Rio Doce-Olinda (c).

**Figure 3.**
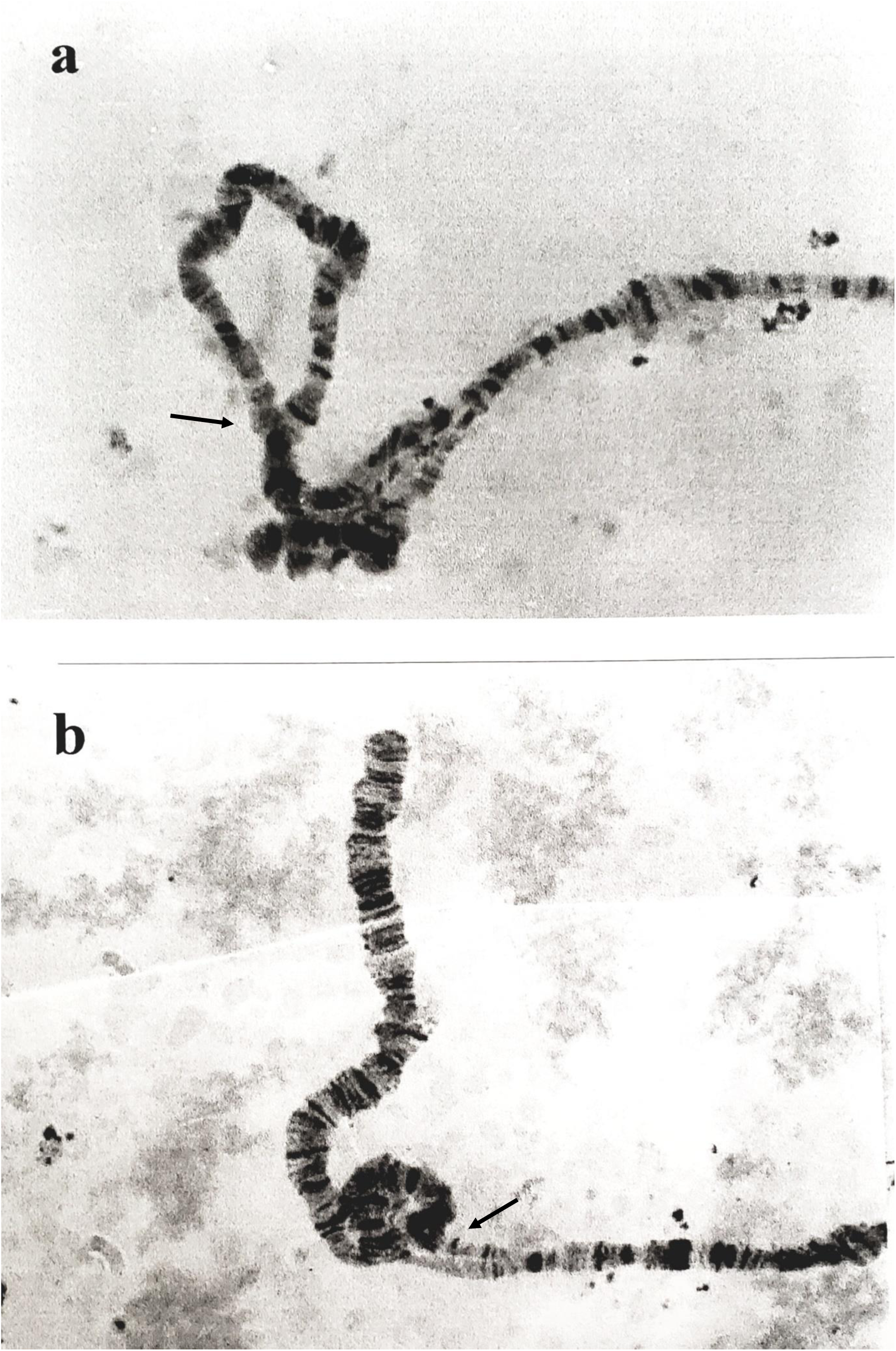
Chromosomal inversions in the IIIL arm of *D. malerkotliana*. In(IIIL)72A-64A, with frequency of 60% of the population of Campus-UFPE (a). In(IIIL)70C-73B, with a frequency of 66% of the Dois Irmãos Park (b).

**Figure 4.**
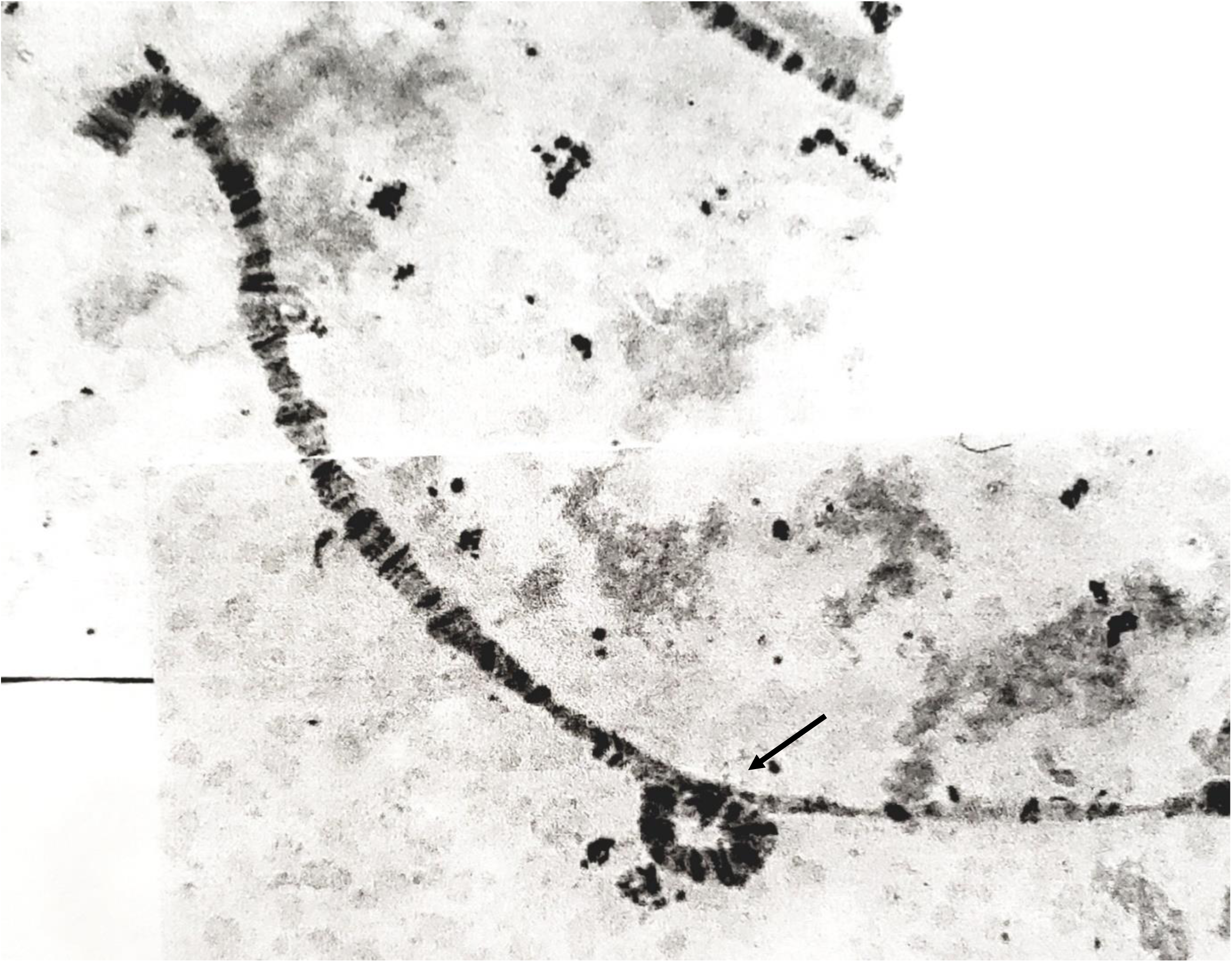
Chromosomal inversion in the IIR-In(IIR)50B-51A arm, with a frequency of 33% in the population of *D. malerkotliana* of Dois irmãos Park.

**Figure 5.**
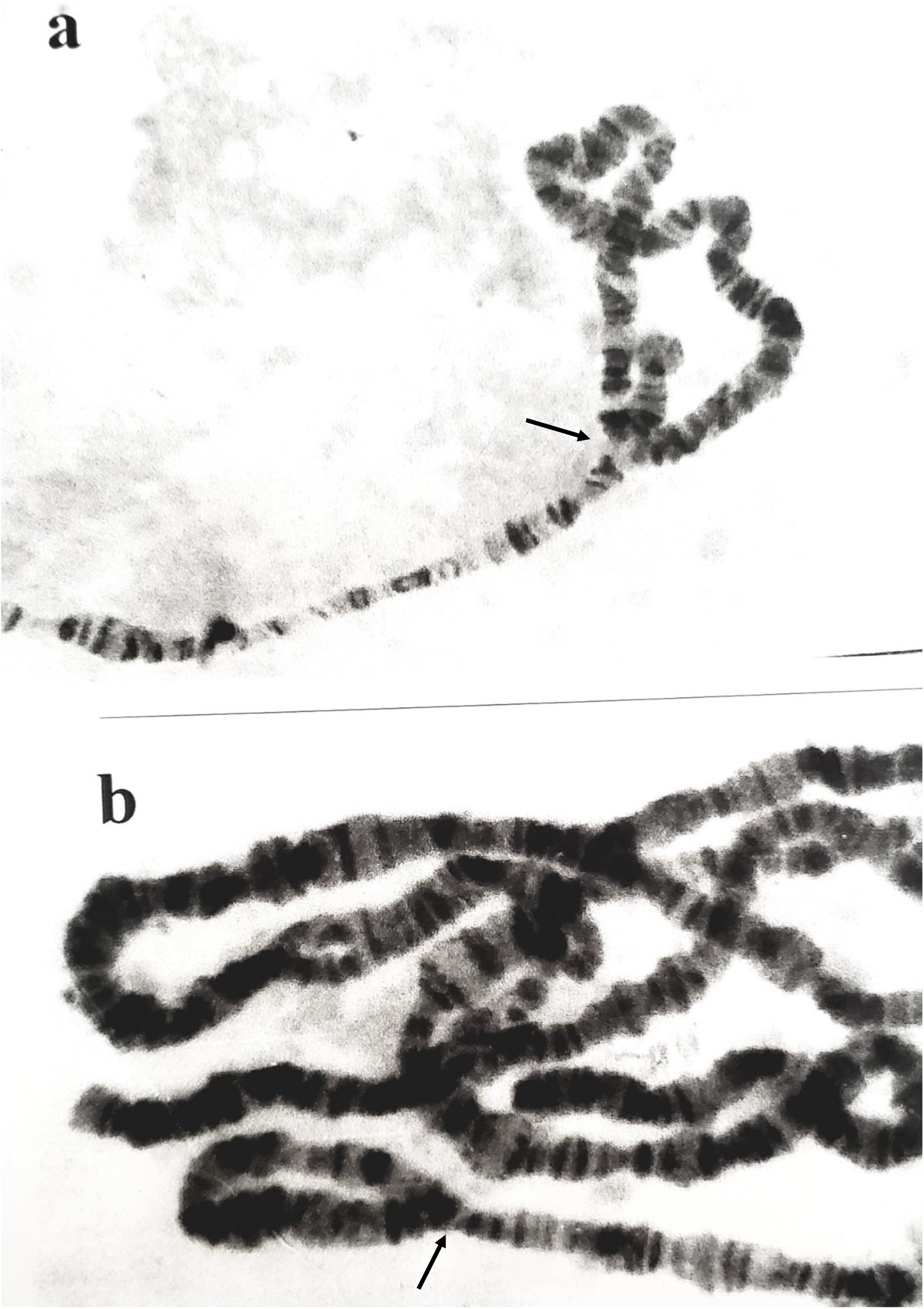
Chromosomal inversion in the IIL arm of *D. malerkotliana*. In (IIL)24B-39A, with frequency of 18.5% of the population of Rio Doce, and with frequency of 33% in the population of Dois Irmãos Park (a). In(IIL)21-25 with a frequency of 100% in the population of Rio Doce (b).

